# Nociception-dependent CCL21 induce dorsal root ganglia axonal growth via CCR7-ERK activation

**DOI:** 10.1101/2022.02.04.479124

**Authors:** Francina Mesquida-Veny, Sara Martinez-Torres, Jose Antonio Del Rio, Arnau Hervera

**Affiliations:** Molecular and Cellular Neurobiotechnology, Institute for Bioengineering of Catalonia (IBEC), Barcelona, Spain; Department of Cell Biology, Physiology and Immunology, University of Barcelona, Barcelona, Spain; Network Centre of Biomedical Research of Neurodegenerative Diseases (CIBERNED), Institute of Health Carlos III, ministry of Economy and Competitiveness, Madrid, Spain; Institute of Neuroscience, University of Barcelona, Barcelona, Spain

**Author notes:** These authors have contributed equally to this work and share first authorship (Francina Mesquida-Veny, Sara Martinez-Torres).

**Keywords:** CCL21, CCR7, MEK-ERK, axonal growth, actin

## Abstract

While chemokines were originally described for their ability to induce cell migration, many studies show how chemokines also take part in a variety of other cell functions, acting as adaptable messengers in the communication between a diversity of cell types. In the nervous system, chemokines participate both in physiological and pathological processes, and while their expression is often described on glial and immune cells, growing evidence describe the expression of chemokines and their receptors in neurons, highlighting, their potential in auto- and paracrine signalling. In this study we analysed the role of nociception in the neuronal chemokinome, and their role in axonal growth. We found that stimulating TRPV1^+^ nociceptors induces a transient increase in CCL21. Interestingly we found that, this CCL21 increases neurite growth of large diameter proprioceptors *in vitro*. Consistent with this, we show that proprioceptors express the CCL21 receptor CCR7, and a CCR7 neutralizing antibody dose-dependently attenuates CCL21-induced neurite outgrowth. Mechanistically, we found that CCL21 binds locally to its receptor CCR7 at the growth cone, activating the downstream MEK-ERK pathway, that in turn activates N-WASP, triggering actin filament ramification in the growth cone, resulting in increased axonal growth.

## 1 Introduction

Classically, chemokines have been associated with leukocyte migration (1). Nevertheless, growing evidence shows they can signal to a great variety of cell types and tissues (2,3). In addition, as conventional chemokine receptors are G-protein coupled receptors (GPCRs), chemokines can initiate a broad variety of intracellular signaling pathways (1).

Several studies have demonstrated the presence and the importance of chemokines in the nervous system (2,4,5). For instance, CXCL12 plays a vital role in regulating neuronal migration during cortical development (6,7). Remarkably, research is now showing neuronal expression of both chemokines and their receptors, suggesting an implication of these cells in direct neuronal communication (5,8)., In homeostasis, neuronal CX_3_CL1 interacts with microglia preventing its activation (9). Additionally, neuronal chemokines have emerged as fundamental signals after insult, as for CCL2 or CCL21 among others (10–13), that are secreted upon neuronal injury and serve as chemoattractants of immune cells, triggering their activation and often leading to neuropathic pain (11–13). However, as in the case of CCL2 after peripheral injury, some chemokines have been also described to promote axonal regeneration as a result of macrophage recruitment and phenotype modulation (10,14). Meanwhile, although several chemokine receptors are expressed in different neuronal types (8,9), little is known about the functions chemokines may exert as autocrine or paracrine messengers on other neurons.

Neurons alter their secretome when exposed to different stimuli and according to their physiological state. In that direction, neuronal activity has shown to modulate neuronal communication, including with microglia or with other neurons beyond classical neurotransmission (15–17). Nociceptor activity after axonal injury is normally associated with pathological neuropathic pain (18), despite that, some studies have started to uncover how nociception participates in the healing process, such as promoting skin or adipose tissue regeneration, as well as neovascularization (19–21). These findings indicate that nociception might function as a key component of the healing machinery, and it is therefore important to study its precise roles in healing and regeneration in different tissues, including the nervous system.

In that sense, injured nociceptors have been described to release CCL21, however, whether this expression affected axonal regeneration has not been previously assessed (13). CCL21 has a primordial role in immune cell homing, via its canonical receptor CCR7 (22), but other functions for this chemokine in distinct tissues are also emerging, such as cartilage regeneration (23) and neuropathic pain induction in the CNS (13). Interestingly, in accordance with its function as a migration cue, the CCL21-CCR7 interaction activates intracellular pathways related to chemotaxis, via ERK signaling, this includes actin cytoskeleton remodeling via RhoA (24). Since cellular migration and growth cone dynamics are analogous mechanisms (25,26), we hypothesized that CCL21 could exert a growth promoting effect on neurons.

In the present study, we investigated the impact of nociceptor activation in the neuronal chemokinome which led us to find an undescribed mechanism of neuronal communication between two different neuronal types, nociceptors and proprioceptors. Specifically, we found that activation of TRPV1^+^ nociceptors induce an increase in CCL21 expression. Moreover, we revealed a novel role for this CCL21 in proprioceptors, promoting neurite outgrowth. We then found that the receptor CCR7, expressed in proprioceptors, was required for this effect, which was also dependent on the MEK-ERK pathway. Finally, our results disclose a local mechanism in the growth cone, where CCR7 expression is concentrated, affecting the actin cytoskeleton, ultimately leading to enhanced neurite outgrowth.

## 2 Materials and methods

### Mice

B6.Cg-Tg(Thy1-COP4/EYFP)18Gfng/J (Thy1-ChR2) (27) were obtained from Jackson Laboratories and C57BL/6J mice from Charles River Laboratories. PV-Cre/Ai27D/CSP-Flox mice (#008069 and #012567 (Jackson Laboratories) crossbreeding) were provided by Dr. Rafael Fernández Chacón (Instituto de Biomedicina de Sevilla, Spain). Mice ranging from 6-10 weeks were used. All animal work was approved by the Ethics Committee on Animal Experimentation (CEEA) of the University of Barcelona (procedures #276/16, 47/20, and OB41/21).

### Compounds

Capsaicin (Merck, 100μg/ml i.pl.), CCL21 (Peprotech, 100ng/sciatic nerve *ex vivo*; 1nM, 10nM, 50nM *in vitro*;), αCCR7 blocking antibody (R&D systems, 2-5μg/ml), rat immunoglobulin (Ig) G (Merck, 2-5μg/ml), U0126 (Promega, 1μM), Wiskostatin (Merck, 1 μM), CCL19 (Peprotech, 1 nM, 10nM, 50nM), Pertussis toxin (Merck, 50ng/ml).

### *In vitro* optogenetic stimulation

Thy1-ChR2 DRG neurons were used for *in vitro* optogenetic stimulation. A 470nm emission LED array (LuxeonRebelTM) under the control of a Driver LED (FemtoBuck, SparkFun) of 600mA and a pulse generator PulsePal (OpenEphys)(28) was used to deliver blue light to neuronal cultures. The optogenetic stimulation protocol consisted in 1h of illumination at 10Hz of frequency with 10ms-90ms pulses, in 1s ON-4s OFF periods. Stimulation was applied 2h after seeding.

### Capsaicin administration

Intraplantar (i.pl.) injection of capsaicin (10μl of 100μg/ml in PBS + 10% ethanol) was performed 2h before animal sacrifice and DRG dissection.

### Dorsal root ganglia (DRG) neuronal culture

DRGs were dissected, collected in ice-cold Hank’s balanced salt solution (HBSS) (ThermoFisher Scientific) and transferred to a digestion solution (5mg/ml Dispase II (Merck) and 2.5mg/ml Collagenase Type II (ThermoFisher Scientific) in DMEM (ThermoFisher Scientific)) incubated for 45min at 37°C. DRGs were then centrifuged and digestion solution was then exchanged for DMEM:F12 (ThermoFisher Scientific) media supplemented with 10% heat-inactivated FBS and 1x B27 (ThermoFisher Scientific), the DRGs were then dissociated by pipetting. The resulting cell suspension was then spun down, resuspended in culture media (DMEM:F-12 media supplemented with 1x B27 and penicillin/streptomycin (P/S) (ThermoFisher Scientific)) and seeded in 48-well plates (3000-4000 cells/well) previously coated with 0.1mg/ml poly-D-lysine (2h, 37°C; Merck) and 2μg/ml laminin (over-night (O/N), RT (room temperature); ThermoFisher Scientific). Cells were kept at 37°C in a 5% CO_2_ atmosphere.

Compounds were added 2h after plating, unless in combinatorial experiments, when blocking antibodies or pharmacological inhibitors were added at 1,5h and the remaining compound 30 min later. Cells were fixed 24h after the treatment.

### Local administration to the sciatic nerve and *ex vivo* DRG culture

Mice were anaesthetized with inhaled isofluorane (4% induction, 2% maintenance). The sciatic nerve was hooked and immobilized after skin incision and blunt dissection of the gluteus maximus and the biceps femoralis. Compounds were locally injected into the nerve and the inscision was closed by layer. Animals were then allowed to recover for 24h. Sciatic DRGs (L4, L5, L6) were dissected and processed for cell culture as previously explained.

### Immunocytochemistry

Cells were fixed with cooled 4% paraformaldehyde (PFA) for 15min and washed in PBS (0.1M). The cells were then blocked (PBS-0.25% Triton X-100+ 1% BSA (bovine serum albumin)) for 1h at RT and then primary antibodies (anti-βIII tubulin (Tuj1, 1:1000, BioLegend), anti-NFH (1:1000, Merck), anti-CCR7 (1:200, Abcam)), were incubated O/N at 4°C in the same solution. After several washes, Alexa-Fluor-conjugated secondary antibodies were incubated in blocking solution for 1h at RT and cell nuclei were counterstained with Hoechst.

### Immunohistochemistry

DRGs were dissected and fixed in PFA 4% for 2h at 4°C. After washing the DRGs were transferred into cryoprotection solution (PBS-30% sucrose) for 24h at 4°C. Tissue freezing medium (OCT, Merck) was used to freeze the DRGs in blocks, and then 10μm slices were generated with a cryostat (Leica CM 1900) and mounted on slides. The slides were washed in TTBS (TBS (Tris-buffered saline) + 1% Tween) and blocking solution (8% BSA and 0.3% Tx and 1/150 mαIgG (Jackson Immuno Research)) was then incubated for 1h at RT. After an O/N incubation with primary antibodies (anti-TRPV/VR1 (1:200, Santa Cruz), anti-CCL21 (1:200 Peprotech), anti-CCR7 (1:200, Abcam)) in 2% BSA and 0.2% Tx in TBS at 4°C, and washing, Alexa-Fluor-conjugated secondary antibodies were added in the same solution for 1h at RT. Preparations were then mounted with coverslips in Mowiol™ (Merck).

### Fluorescence intensity analysis

DRG slices were stained for CCL21 and TRPV1. Images were acquired with a LSM 800 confocal microscope (Zeiss) using an AxioCam 503c camera (Zeiss) at 20X magnification. Mean CCL21 fluorescence intensity (MFI) was computed in TRPV1^+^ cells using ImageJ, subtracting the background for each image.

### Neurite length analysis

Tuj1 immunocytochemistry was performed and imaged at 10X magnification using an Olympus microscope IX71 with an Orca Flash 4 (3 images per well). The neurite length of large-diameter (>35μm) neurons was blindly quantified with the Neuron J plugin for ImageJ (29). Average neurite length per neuron was computed.

### Statistical analysis

Statistics and graphical representation were carried out using Prism 6.0 (GraphPad™ Software). Shapiro-Wilk test was used to verify normality of the distributions. * or # indicate significant differences in ANOVA followed by Bonferroni’s post-hoc test or Student’s t-test. Plotted data represents mean±s.e.m (standard error of the mean). All tests performed were two-sided, and adjustments for multiple comparisons and/or significantly different variances (Fisher’s F) were applied were indicated. All data analysis was performed blind to the experimental group by two independent experimenters. Unless otherwise stated, sample size was chosen in order to ensure a power of at least 0.8, with a type I error threshold of 0.05, in view of the minimum effect size that was expected.

## 3 Results

### 3.1 CCL21 expression is upregulated upon nociceptor activation

While most defined neuronal chemokines are induced after traumatic injury or inflammatory signalling (30), we analysed whether stimulating neuronal activity in the DRG would have an impact in chemokine expression and secretion. To this aim we used DRG neuronal cultures expressing ChR2 (from Thy1-ChR2 animals) and subjected them to optical stimulation. We recovered the media 24h later and measured chemokine secretion using a Mouse Chemokine Array (Raybiotech). Interestingly, we found a remarkable increase in CCL21 levels compared to non-stimulated controls (Data not shown). ChR2 expression in DRGs is ubiquitous among all sensory subtypes (Fig 1A-B), however, previous studies had reported CCL21 expression specifically in small diameter TRPV1^+^ nociceptors after peripheral nerve injury (31,32), so we hypothesized that this CCL21 increase after optogenetic stimulation could be specific of this neuronal subtype. Accordingly, CCL21 expression was increased in TRPV1^+^ nociceptors 2h after i.pl capsaicin (TRPV1 agonist) injection as compared to vehicle administered animals (Student’s T test *p*= 0.0347) (Fig. 1C-E).

**Figure 1.**
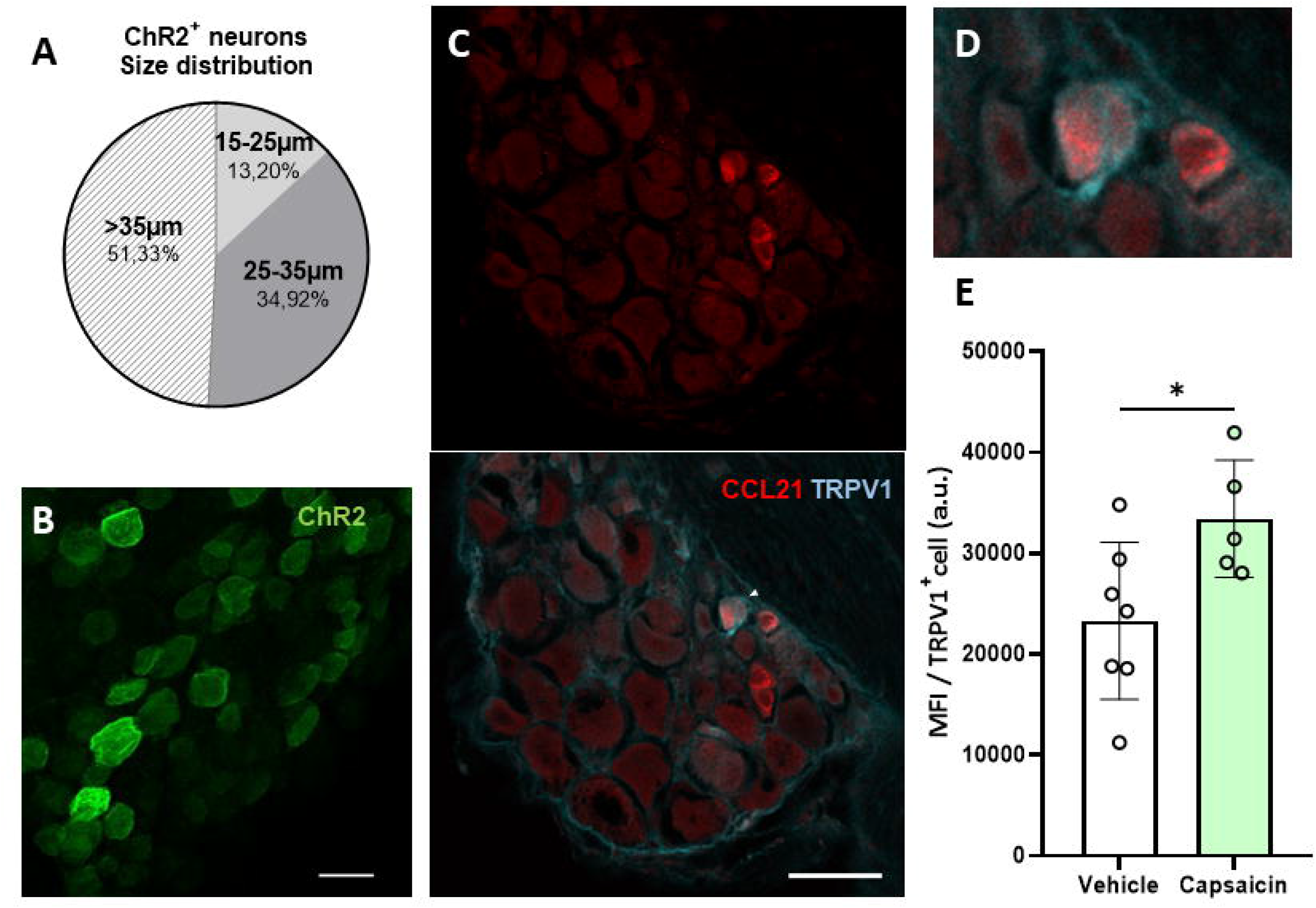
CCL21 is upregulated upon TRPV1^+^ nociceptive neuron activation. A. Graph showing the proportion of ChR2^+^ cells depending on the neuronal diameter. ChR2 is expressed in both large-diameter (>35μm) and small-diameter (35-15μm) neurons. B. ChR2 expression in Thy1-ChR2 DRGs. Scale bar: 50μm. C. Immunohistochemistry showing CCL21 expression in TRPV1^+^ neurons 2h after capsaicin injection. White arrows: magnified neurons in D. Scale bar: 50μm. D. High magnification inset. E. Intraplantar capsaicin injection increased CCL21 expression in TRPV1^+^ nociceptors specifically as shown by mean fluorescence intensity (MFI) of TRPV1^+^ cells. a.u.: arbitrary units. Data are expressed as mean±s.e.m; Student’s t-test; \**p* < 0.05; n=5-7 images.

### 3.2 CCL21 promotes neurite outgrowth

To test whether this chemokine could also influence axonal growth, we administered CCL21 into the sciatic nerve and cultured disaggregated DRG neurons 24h after. This led to an increment in the neurite outgrowth of *ex vivo* CCL21-treated DRG neurons when compared to vehicle treated ones (Student’s t-test p= 0.0046) (Fig. 2A-B). Additionally, administration of CCL21 at different doses (1nM, 10nM, 50nM) on DRG cultures, resulted in a dose-dependent increase of DRG neurite outgrowth *in vitro* (ANOVA followed by Bonferroni test; Veh vs 50nM *p*= 0.0129; 1nM vs 50nM *p*= 0.0440) (Fig. 2C-D).

**Figure 2.**
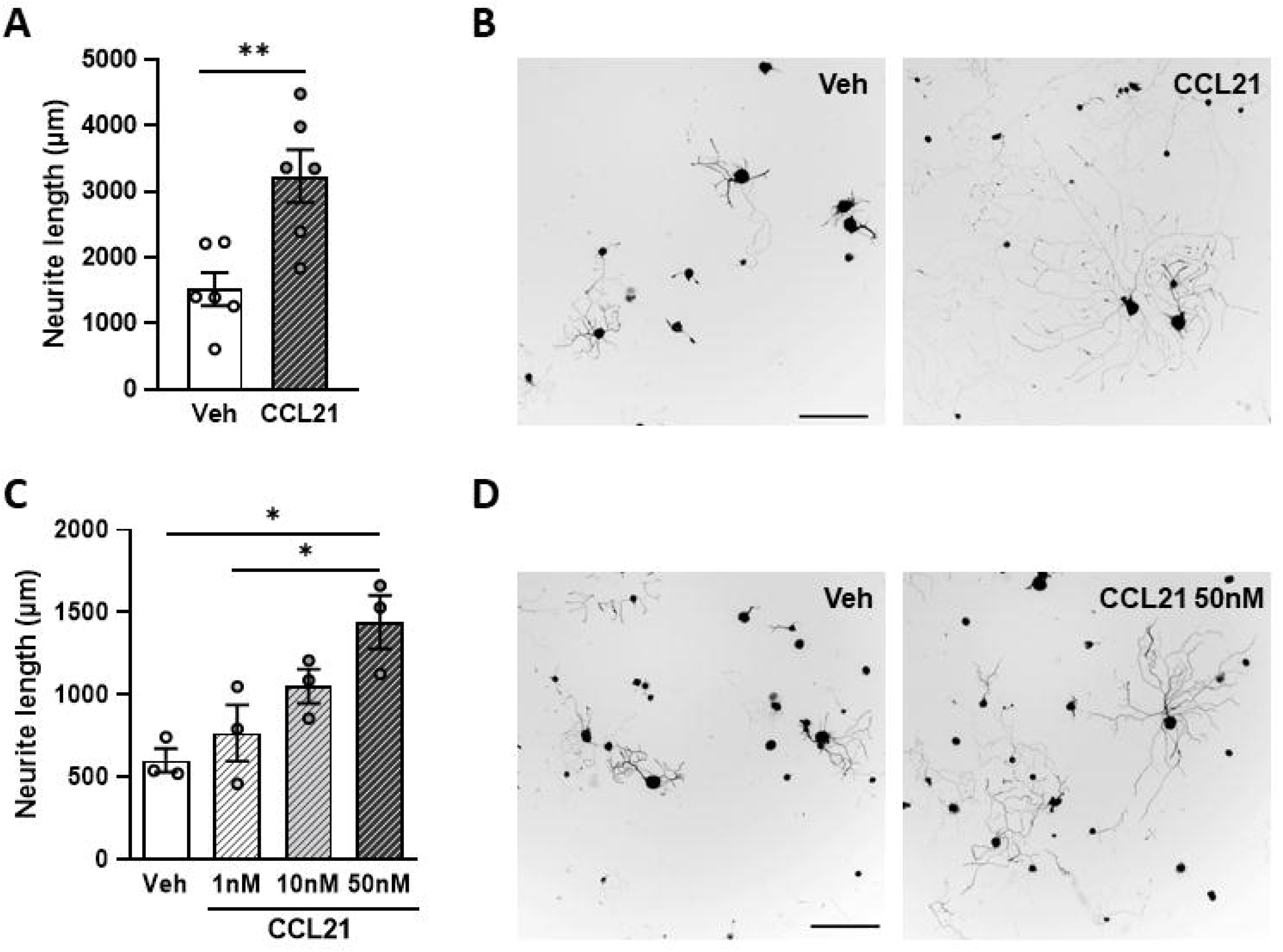
CCL21 enhances DRG neurite outgrowth. A. *Ex vivo* CCL21 administration in the sciatic nerve promoted neurite outgrowth of DRG large-diameter neurons. Data are expressed as average neurite length per neuron±s.e.m; Student’s t-test; \*\**p* < 0.01; n=6 sciatic nerves. C. Dose-dependent increase in neurite length of *in vitro* CCL21-treated DRG large-diameter neurons. Data are expressed as average neurite length per neuron±s.e.m; One-way ANOVA, Bonferroni’s post-hoc; \**p* < 0.05; n=3 wells. B-D. Tuj-1 representative immunostainings (B: *ex vivo* D: *in vitro*) after 24h in culture. Scale bar: 250μm.

### 3.3 CCL21 activates proprioceptive CCR7 to promote neurite outgrowth

CCL21 is a functional ligand of CCR7 (33). We therefore analysed the expression pattern of CCR7 in the DRG. We found CCR7 expression mainly in neurons, including in parvalbumin^+^ (PV) neurons, that correspond to proprioceptors (Fig. 3A). Additionally, administering a CCR7-blocking antibody to DRG cultures inhibited the neurite outgrowth induced by CCL21 as compared to the IgG control (two-way ANOVA followed by Bonferroni test; interaction p= 0.0233; IgG-veh vs IgG-CCL21 p= 0.0055; IgG-CCL21 vs αCCR7 2μg-veh p= 0.0040; IgG-CCL21 vs αCCR7 2μg-CCL21 p= 0.0355; IgG-CCL21 vs αCCR7 5μg-veh p= 0.0037; IgG-CCL21 vs αCCR7 5μg-CCL21 p= 0.0079) (Fig. 3B-C). Parallelly, we tested whether CCL19, another CCR7 ligand(33), would induce the same effects. Oppositely, CCL19 did not result in increased neurite outgrowth when administered *in vitro* (Kruskal-Wallis test followed by Dunn’s test; Veh vs 50nM p > 0.9999) (Supplementary Figure 1), suggesting a CCL21 biased CCR7 activation is responsible for this particular mechanism.

**Figure 3.**
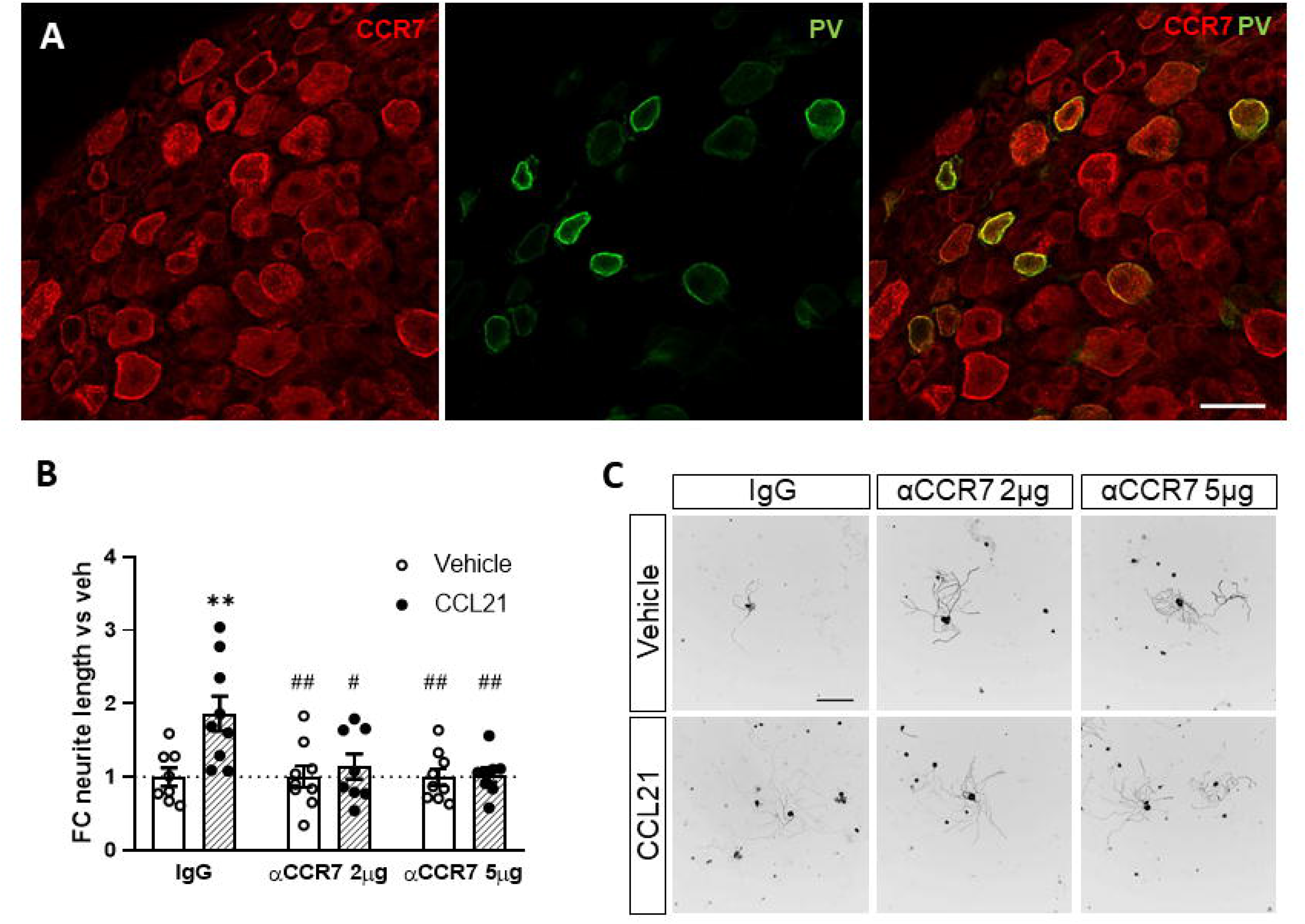
CCR7 is required for CCL21-mediated DRG outgrowth. A. PV^+^ neurons express the canonical CCL21 receptor CCR7. Scale bar: 50μm. B. CCR7-blockade abolished the CCL21-dependent growth induction. Data are expressed as mean fold change of average neurite length per neuron vs each vehicle group±s.e.m. Two-way ANOVA, Bonferroni’s post-hoc; \*\**p* < 0.01 (vs IgG-veh); #*p* < 0.05; ##*p* < 0.01 (vs IgG-CCL21); n=8-9 images. C. Tuj-1 representative immunostainings after 24h in culture. Scale bar: 250μm.

### 3.4 CCL21-CCR7 activates MEK-ERK pathway

We then targeted the two main known downstream actuators of CCR7 activation, the MEK pathway and the G_i/o_ protein previously known to be involved in axon growth(34–36)). Pharmacological inhibition of MEK with U0126 blocks the neurite outgrowth induced by CCL21 (Fig. 4A-B), as evidenced by the significant interaction of the treatments (*p*= 0.0117) on the two-way ANOVA (multiple comparisons with Bonferroni test: DMSO-Veh vs DMSO-CCL21 *p*= 0.0005; DMSO-CCL21 vs U0126-Veh *p*= 0.0004; DMSO-CCL21 vs U0126-CCL21 *p*= 0.0001). Conversely, pertussis toxin (Ptx) administration, an inhibitor of the Gi/o protein did not affect the CCL21-induced outgrowth (Supplementary Figure 2) as evidenced by the lack of significant interaction of the treatments (*p*= 0.7961) on the two-way ANOVA (Student’s t-test w/o (without) Ptx: *p*= 0.0115; with Ptx: *p*= 0.7458).

**Figure 4.**
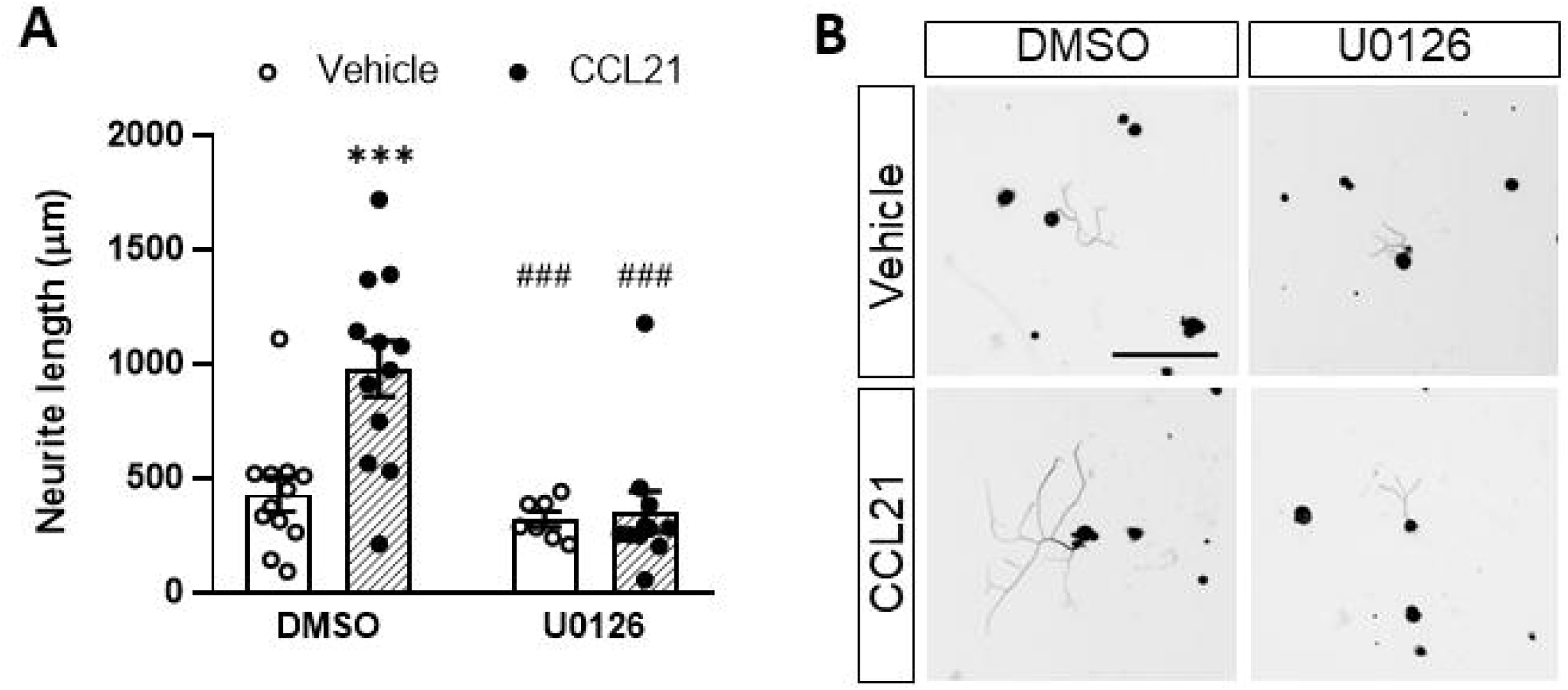
Inhibiting the MEK-ERK pathway impairs the CCL21-dependent increased outgrowth. A. CCL21 and U0126, a MEK inhibitor, co-administration resulted in reduced neurite outgrowth compared to CCL21 administration alone. Data are expressed as average neurite length per neuron±s.e.m; Two-way ANOVA, Bonferroni’s post-hoc; \*\*\**p* < 0.001 (vs veh-DMSO), ###p < 0.001 (vs CCL21-DMSO). n=7-12 images. B. Tuj-1 representative immunostainings after 24h in culture. Scale bar: 250μm.

### 3.5 CCL21-CCR7-MEK pathway modulates actin cytoskeleton to promote neurite outgrowth

Local assessment of the axonal tips revealed larger growth cones after CCL21 treatment (Fig. 5A), this is in consonance with the especially abundant CCR7 expression found on these structures (Fig. 5B). We then sought to check the local effects that CCL21 could have in cytoskeletal dynamics of the growth cones. Consequently, we inhibited actin branching by combining CCL21 administration with wiskostatin, an inhibitor of the neural Wiskott-Aldrich syndrome protein (N-WASP), that acts as an Arp2/3 complex activator. Wiskostatin co-administration greatly reduced the CCL21-induced growth, supporting a local effect of CCL21 in the growth cone (two-way ANOVA followed by Bonferroni test; interaction p= 0.0825; one-way ANOVA followed by Bonferroni test; DMSO-Veh vs DMSO-CCL21 p= 0.0207; DMSO-CCL21 vs Wiskostatin-Veh p= 0.0202).

**Figure 5.**
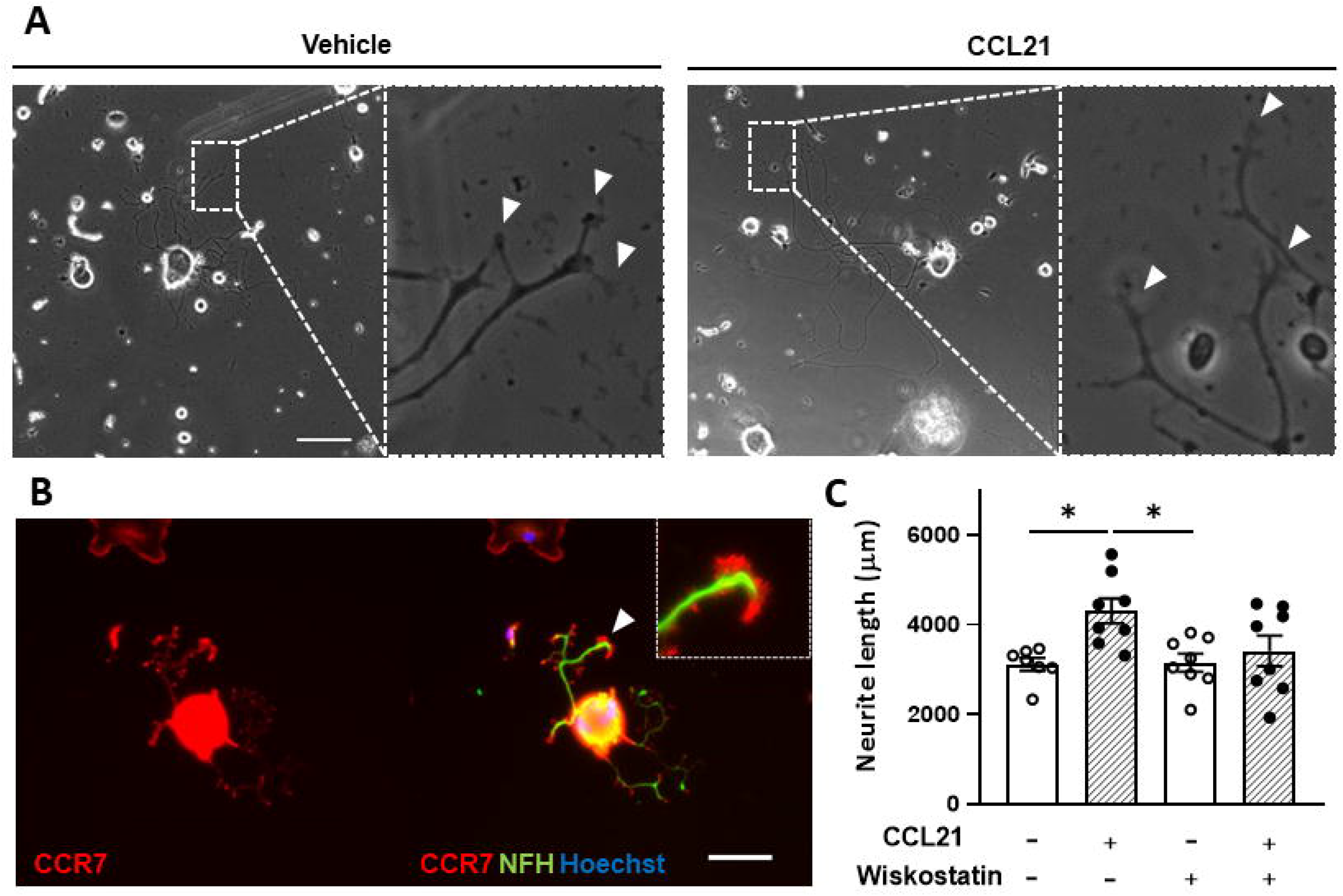
CCL21 stimulates actin dynamics in the growth cone. A. CCL21 administration resulted in enlarged growth cones in large-diameter DRG neurons. White arrows: growth cones. Scale bar: 100μm. B. CCR7 expression is specially elevated in the growth cone. Scale bar: 50μm. C. Actin branching is important for CCL21-dependent neurite outgrowth, as shown by impaired growth after wiskostatin treatment. Data are expressed as average neurite length per neuron±s.e.m; One-way ANOVA, Bonferroni’s post-hoc; \**p* < 0.05; n=7-8 wells.

## 4 Discussion

There are plenty of evidence that neuronal activity can alter neuronal signalling. In that sense, we found that the expression and release of the chemokine CCL21 is enhanced upon neuronal stimulation. Interestingly, we found that specific TRPV1^+^ nociceptor stimulation is responsible for CCL21 production, similarly to what has already been described upon axonal injury (31).

After axonal injury, release of neuronal CCL21 is mainly linked to neuropathic pain (13,37), however, its role in axon growth and regeneration had never been described before. Interestingly, our results revealed that CCL21 induces growth in proprioceptive neurons trough activation of the CCR7-MEK-ERK pathway, exerting an effect on actin dynamics at the growth cone level.

CCL21 was first designated as a recruiting cue for leucocytes, specifically stimulating the migration of T cell subpopulations and dendritic cells (22,38–40). More recently, it has also been described to induce migration in other cells such as tumorigenic or mesenchymal stem cells (41–45). Fundamentally, cell migration and axonal growth are events that share similar cellular and molecular machinery (46,47). Thus, molecules that orchestrate one process will most likely be implicated in the other, and vice versa, as for instance what occurs with CXCL12 (48–51).

We also describe that CCL21 executes its growth-inducing function through its canonical receptor CCR7, in accordance, we found abundant expression of this receptor on proprioceptive (PV^+^) neurons, similarly to the findings of other studies, where neuronal CCR7 is abundantly found in peripheral nerves and hippocampal neurons (52,53). Activation of CCR7 has already been shown to play a cardinal role in the CCL21-induced cell migration (42,44,47,54,55), that, as stated, activates similar cellular and molecular machinery than axonal growth (46,47).

Intriguingly, CCL19, another CCR7 ligand, did not induce axonal growth. This finding strengthens the view of biased ligand-receptor responses, as already shown for CCL21 and CCL19, which can trigger biased downstream effectors of CCR7 (56,57). These biased activations result in particular mechanisms activated only by CCL21; however, this effect often varies depending on the target cell (58–61).

We also found that pharmacological inhibition of the MEK-ERK pathway, one of the main downstream mediators of CCR7 effects, inhibited the growth induced by CCL21. This goes in line other studies defining MEK-ERK as the underlying mechanism of the chemotaxis induced by CCL21-CCR7 (55,62). Contrarily, we did not see an effect by inhibiting the G_i/o_ protein in the CCL21-induced outgrowth, however lack of growth-suppression could derive from inactivation of other growth-inhibitor pathways activated by the G_i/o_ (63).

MEK-ERK pathway activation has been previously implicated in axonal regeneration (64–66). For instance, after peripheral conditioning injury, ERK is phosphorylated and has been shown to affect multiple cellular processes affecting axonal growth, including transcriptional and epigenetic alterations, resulting increased expression of several regeneration associated genes (RAGs) (66), increased retrograde transport (64,67) as well as stimulation of cytoskeleton dynamics (65). According with the latter, ERK has been shown to have a direct effect on actin polymerization (68,69), through phosphorylation of different effectors such as WAVE2, cortactin and Rac1 (70–74). Remarkably, we found CCR7 expression to be particularly elevated in the growth cones, therefore, we evaluated the effects of actin dynamics in the growth cone as a putative mechanism of the CCL21-induced growth. In that sense, coadministration of an N-WASP pharmacological inhibitor, a key actin polymerization component, blocked the growth induced by CCL21. N-WASP is an Arp2/3 complex activator, which in turn works as an actin-binding protein, mainly responsible for actin filament branching (75). Previous studies already showed the essential role of Arp2/3 in growth cone progression (76,77). Mechanistically, ERK is known to phosphorylate cortactin, leading to N-WASP binding and activation (71). Thus, in a nutshell, we found that upon CCL21 binding to its receptor CCR7 in the axonal growth cone, there is a downstream MEK-ERK activation, that leads to N-WASP activation, triggering actin filament branching in this structure, resulting in increased axonal growth (Figure 6).

**Figure 6.**
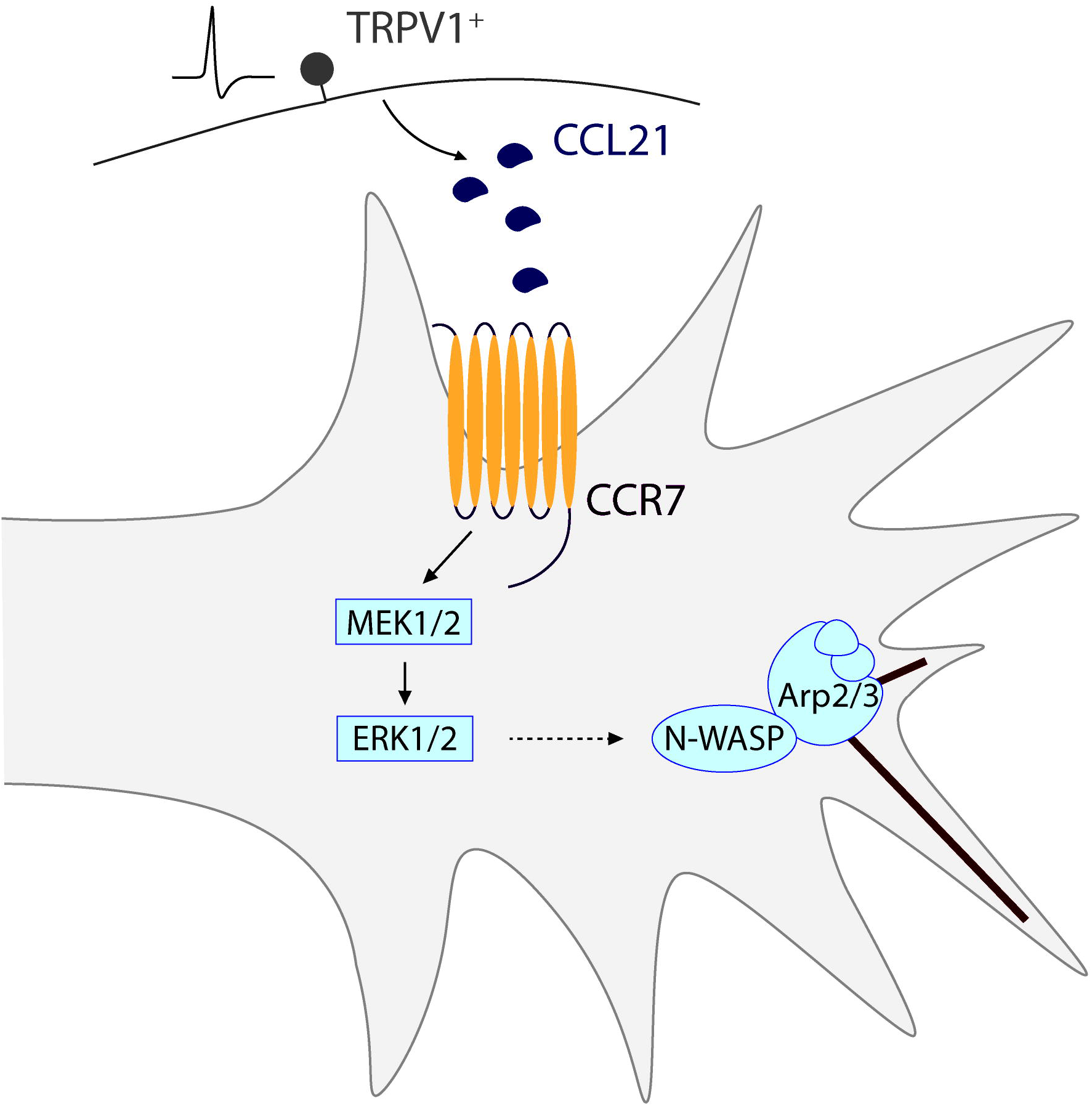
Schematic representation of the proposed mechanism for a novel nociceptor-proprioceptor dialogue leading to neuritic growth. Activated TRPV1+ nociceptors secrete CCL21 which promotes actin branching in the growth cone of proprioceptor neurons. This mechanism is mediated by the CCL21-CCR7 interaction, leading to a downstream activation of the MEK-ERK pathway and final N-WASP-related actin cytoskeleton modifications.

While CCL21 has been already described for its role in regeneration in other tissues including skin, cartilage, and vascular remodelling (41,43,78), here we describe a novel role of this chemokine in axonal growth.

Additionally, the mechanism described involves an unprecedented paracrine dialogue based on chemokines between two different DRG neuronal types. While neurons are highly interconnected through synapses, there is little work regarding other forms of neuronal communication (79). Growth factors, for instance, have been implicated in both paracrine and autocrine neuronal communication (80–83), however, we describe a novel chemokine dependant mechanism of neuron-neuron signalling.

Parallelly, we have also unravelled a novel role of nociceptive signalling in sensory axon regeneration. Nociception and pain are tightly associated with tissue injury (84,85), and while it is not surprising that a signal initiated by an injury could trigger the healing process, after nerve injury chemokine release and nociceptor activity have been typically linked to pathological neuropathic pain. Contrarily, here we show that stimulating nociceptor activity triggers the secretion of a growth-promoting chemokine, CCL21. In agreement with this, nociceptive signalling participates in the healing cascade of several tissues, for example, nociceptor activation induces adipose tissue regeneration through CGRP (calcitonin gene-related peptide) secretion (86), angiogenesis via Substance-P-mediated effects (87), and skin regeneration through modulation of the immune response (88). Pain and nociceptive signalling are complex evolutionary mechanisms that might have further implications than previously anticipated, as they play central roles orchestrating and promoting the healing process in different tissues.

This also suggest caution in the indiscriminate use of analgesic drugs and treatments after injury, as these may hinder nociceptive regenerative signalling, limiting the healing process, similarly to the effects observed by broad immunosuppressive drugs (89) or antioxidants (90) after SCIs. Therefore, appropriate timing and level of analgesic inhibition after injury will likely need to be tailored to provide pain relief while avoiding unwanted effects in hindering the tissue regeneration. An additional intriguing implication is the possibility to modulate nociceptive signalling to achieve tissue regeneration. While therapeutically, inducing nociception is not a reasonable approach, further investigation and characterization of signalling elicited by nociceptor stimulation may increase our understanding of the molecular mechanisms underlying the healing process, and may enable the future design of therapeutic targets and strategies to foster tissue regeneration.

## 5 Conflict of Interest

The authors declare that the research was conducted in the absence of any commercial or financial relationships that could be construed as a potential conflict of interest.

## 6 Author Contributions

FMV performed, designed experiments, performed data analysis, and wrote the manuscript; SMT performed and designed experiments and performed data analysis; JADR supervised experiments, provided experimental funds and edited the manuscript; AH performed, designed experiments, performed data analysis, provided experimental funds and wrote the manuscript.

## 7 Funding

This research was supported by HDAC3-EAE-SCI Project with ref. PID2020-119769RA-I00 from MCIN/ AEI /10.13039/501100011033 to AH and PRPSEM Project with ref. RTI2018-099773-B-I00 from MCINN/AEI/10.13039/501100011033/ FEDER “Una manera de hacer Europa”, the CERCA Programme, and the Commission for Universities and Research of the Department of Innovation, Universities, and Enterprise of the Generalitat de Catalunya (SGR2017-648) to JADR. The project leading to these results received funding from the María de Maeztu Unit of Excellence (Institute of Neurosciences, University of Barcelona) MDM-2017-0729 and Severo Ochoa Unit of Excellence (Institute of Bioengineering of Catalonia) CEX2018-000789-S from MCIN/AEI/10.13039/501100011033. FMV was supported by a fellowship from the “Ayudas para la Formación de Profesorado Universitario” (FPU16/03992) program, from the Spanish Ministry of Universities.

## 8 Acknowledgments

The authors thank Miriam Segura-Feliu for her technical help. PV-Cre/Ai27D/CSP-Flox mice (#008069 and #012567 (Jackson Laboratories) crossbreeding) were generated and provided by Dr. Rafael Fernández-Chacón (Instituto de Biomedicina de Sevilla).

## 10 Figure Legends

**Suppl. Fig. 1**. **CCL19 administration does not induce neurite outgrowth.** Data are expressed as average neurite length per neuron±s.e.m; n=8-12 images

**Suppl. Fig. 2**. **Pertussis toxin addition does not prevent the CCL21-dependent neurite outgrowth increase.** Data are expressed as mean fold change of average neurite length per neuron vs each vehicle ±s.e.m; n=9-12 images.

## Notes

### Competing Interest Statement

The authors have declared no competing interest.

## References

1. Hughes CE, Nibbs RJB. A guide to chemokines and their receptors. FEBS Journal (2018) 285:2944–2971. doi: 10.1111/febs.14466

2. Jaerve A, Müller HW. Chemokines in CNS injury and repair. Cell and Tissue Research (2012) doi: 10.1007/s00441-012-1427-3

3. Anders HJ, Romagnani P, Mantovani A. Pathomechanisms: Homeostatic chemokines in health, tissue regeneration, and progressive diseases. Trends in Molecular Medicine (2014) 20:154–165. doi: 10.1016/j.molmed.2013.12.002

4. De Haas AH, Van Weering HRJ, De Jong EK, Boddeke HWGM, Biber KPH. Neuronal chemokines: Versatile messengers in central nervous system cell interaction. Molecular Neurobiology (2007) doi: 10.1007/s12035-007-0036-8

5. Mesquida-Veny F, Del Río JA, Hervera A. Macrophagic and microglial complexity after neuronal injury. Progress in Neurobiology (2021) 200: doi: 10.1016/j.pneurobio.2020.101970

6. Borrell V, Marín O. Meninges control tangential migration of hem-derived Cajal-Retzius cells via CXCL12/CXCR4 signaling. Nature Neuroscience (2006) 9:1284–1293. doi: 10.1038/nn1764

7. Lysko DE, Putt M, Golden JA. SDF1 regulates leading process branching and speed of migrating interneurons. Journal of Neuroscience (2011) 31:1739–1745. doi: 10.1523/JNEUROSCI.3118-10.2011

8. Cartier L, Hartley O, Dubois-Dauphin M, Krause KH. Chemokine receptors in the central nervous system: Role in brain inflammation and neurodegenerative diseases. Brain Research Reviews (2005) 48:16–42. doi: 10.1016/j.brainresrev.2004.07.021

9. Williams JL, Holman DW, Klein RS. Chemokines in the balance: Maintenance of homeostasis and protection at CNS barriers. Frontiers in Cellular Neuroscience (2014) doi: 10.3389/fncel.2014.00154

10. Kwon MJ, Shin HY, Cui Y, Kim H, Le Thi AH, Choi JY, Kim EY, Hwang DH, Kim BG. CCL2 mediates neuron-macrophage interactions to drive proregenerative macrophage activation following preconditioning injury. Journal of Neuroscience (2015) doi: 10.1523/JNEUROSCI.1924-15.2015

11. Hu Z, Deng N, Liu K, Zeng W. DLK mediates the neuronal intrinsic immune response and regulates glial reaction and neuropathic pain. Experimental Neurology (2019) doi: 10.1016/j.expneurol.2019.113056

12. Wang Q, Zhang S, Liu T, Wang H, Liu K, Wang Q, Zeng W. Sarm1/Myd88-5 Regulates Neuronal Intrinsic Immune Response to Traumatic Axonal Injuries. Cell Reports (2018) doi: 10.1016/j.celrep.2018.03.071

13. Biber K, Tsuda M, Tozaki-Saitoh H, Tsukamoto K, Toyomitsu E, Masuda T, Boddeke H, Inoue K. Neuronal CCL21 up-regulates microglia P2X4 expression and initiates neuropathic pain development. EMBO Journal (2011) doi: 10.1038/emboj.2011.89

14. Bhardwaj D, Náger M, Camats J, David M, Benguria A, Dopazo A, Cantí C, Herreros J. Chemokines induce axon outgrowth downstream of hepatocyte growth factor and TCF/ß-catenin signaling. Frontiers in Cellular Neuroscience (2013) doi: 10.3389/fncel.2013.00052

15. Park S, Koppes RA, Froriep UP, Jia X, Achyuta AKH, McLaughlin BL, Anikeeva P. Optogenetic control of nerve growth. Scientific Reports (2015) 5:1–9. doi: 10.1038/srep09669

16. Ferro A, Auguste YSS, Cheadle L. Microglia, Cytokines, and Neural Activity: Unexpected Interactions in Brain Development and Function. Frontiers in Immunology (2021) 12:1–14. doi: 10.3389/fimmu.2021.703527

17. Nguyen PT, Dorman LC, Pan S, Vainchtein ID, Han RT, Nakao-Inoue H, Taloma SE, Barron JJ, Molofsky AB, Kheirbek MA, et al. Microglial Remodeling of the Extracellular Matrix Promotes Synapse Plasticity. Cell (2020) 182:388–403.e15. doi: 10.1016/j.cell.2020.05.050

18. Costigan M, Scholz J, Woolf CJ. Neuropathic pain: A maladaptive response of the nervous system to damage. Annual Review of Neuroscience (2009) 32:1–32. doi: 10.1146/annurev.neuro.051508.135531

19. Amadesi S, Reni C, Katare R, Meloni M, Oikawa A, Beltrami AP, Avolio E, Cesselli D, Fortunato O, Spinetti G, et al. Role for substance P-based nociceptive signaling in progenitor cell activation and angiogenesis during ischemia in mice and in human subjects. Circulation (2012) 125:1774–1786. doi: 10.1161/CIRCULATIONAHA.111.089763

20. Rabiller L, Labit E, Guissard C, Gilardi S, Guiard BP, Moulédous L, Silva M, Mithieux G, Pénicaud L, Lorsignol A, et al. Pain sensing neurons promote tissue regeneration in adult mice. npj Regenerative Medicine (2021) 6:1–9. doi: 10.1038/s41536-021-00175-7

21. Wei JJ, Kim HS, Spencer CA, Brennan-Crispi D, Zheng Y, Johnson NM, Rosenbach M, Miller C, Leung DH, Cotsarelis G, et al. Activation of TRPA1 nociceptor promotes systemic adult mammalian skin regeneration. Science Immunology (2020) 5:1–9. doi: 10.1126/SCIIMMUNOL.ABA5683

22. Förster R, Davalos-Misslitz AC, Rot A. CCR7 and its ligands: Balancing immunity and tolerance. Nature Reviews Immunology (2008) 8:362–371. doi: 10.1038/nri2297

23. Joutoku Z, Onodera T, Matsuoka M, Homan K, Momma D, Baba R, Hontani K, Hamasaki M, Matsubara S, Hishimura R, et al. CCL21/CCR7 axis regulating juvenile cartilage repair can enhance cartilage healing in adults. Scientific Reports (2019) 9:1–12. doi: 10.1038/s41598-019-41621-3

24. Rodríguez-Fernández JL, Criado-García O. The Chemokine Receptor CCR7 Uses Distinct Signaling Modules With Biased Functionality to Regulate Dendritic Cells. Frontiers in Immunology (2020) 11:1–10. doi: 10.3389/fimmu.2020.00528

25. Aberle H. Axon guidance and collective cell migration by substrate-derived attractants. Frontiers in Molecular Neuroscience (2019) 12:1–7. doi: 10.3389/fnmol.2019.00148

26. Cooper JA. Mechanisms of cell migration in the nervous system. Journal of Cell Biology (2013) 202:725–734. doi: 10.1083/jcb.201305021

27. Arenkiel BR, Peca J, Davison IG, Feliciano C, Deisseroth K, Augustine GJJ, Ehlers MD, Feng G. In Vivo Light-Induced Activation of Neural Circuitry in Transgenic Mice Expressing Channelrhodopsin-2. Neuron (2007) 54: doi: 10.1016/j.neuron.2007.03.005

28. Siegle JH, López AC, Patel YA, Abramov K, Ohayon S, Voigts J. Open Ephys: An open-source, plugin-based platform for multichannel electrophysiology. Journal of Neural Engineering (2017) 14: doi: 10.1088/1741-2552/aa5eea

29. Meijering E, Jacob M, Sarria JCF, Steiner P, Hirling H, Unser M. Design and Validation of a Tool for Neurite Tracing and Analysis in Fluorescence Microscopy Images. Cytometry Part A (2004) 58: doi: 10.1002/cyto.a.20022

30. Mesquida-Veny F, del Río JA, Hervera A. Macrophagic and microglial complexity after neuronal injury. Progress in Neurobiology (2021) 200: doi: 10.1016/j.pneurobio.2020.101970

31. Biber K, Tsuda M, Tozaki-Saitoh H, Tsukamoto K, Toyomitsu E, Masuda T, Boddeke H, Inoue K. Neuronal CCL21 up-regulates microglia P2X4 expression and initiates neuropathic pain development. EMBO Journal (2011) doi: 10.1038/emboj.2011.89

32. Zhao P, Waxman SG, Hains BC. Modulation of thalamic nociceptive processing after spinal cord injury through remote activation of thalamic microglia by cysteine-cysteine chemokine ligand 21. Journal of Neuroscience (2007) doi: 10.1523/JNEUROSCI.2209-07.2007

33. Yoshida R, Nagira M, Kitaura M, Imagawa N, Imai T, Yoshie O. Secondary lymphoid-tissue chemokine is a functional ligand for the CC chemokine receptor CCR7. Journal of Biological Chemistry (1998) 273: doi: 10.1074/jbc.273.12.7118

34. Hasegawa Y, Fujitani M, Hata K, Tohyama M, Yamagishi S, Yamashita T. Promotion of axon regeneration by myelin-associated glycoprotein and Nogo through divergent signals downstream of Gi/G. Journal of Neuroscience (2004) 24:6826–6832. doi: 10.1523/JNEUROSCI.1856-04.2004

35. Perlson E, Hanz S, Ben-Yaakov K, Segal-Ruder Y, Seger R, Fainzilber M. Vimentin-dependent spatial translocation of an activated MAP kinase in injured nerve. Neuron (2005) 45:715–726. doi: 10.1016/j.neuron.2005.01.023

36. Chierzi S, Ratto GM, Verma P, Fawcett JW. The ability of axons to regenerate their growth cones depends on axonal type and age, and is regulated by calcium, cAMP and ERK. European Journal of Neuroscience (2005) 21:2051–2062. doi: 10.1111/j.1460-9568.2005.04066.x

37. De Jong EK, Dijkstra IM, Hensens M, Brouwer N, Van Amerongen M, Liem RSB, Boddeke HWGM, Biber K. Vesicle-mediated transport and release of CCL21 in endangered neurons: A possible explanation for microglia activation remote from a primary lesion. Journal of Neuroscience (2005) doi: 10.1523/JNEUROSCI.1019-05.2005

38. Shannon LA, Calloway PA, Welch TP, Vines CM. CCR7/CCL21 migration on fibronectin is mediated by phospholipase Cγ1 and ERK1/2 in primary T lymphocytes. Journal of Biological Chemistry (2010) 285:38781–38787. doi: 10.1074/jbc.M110.152173

39. Britschgi MR, Favre S, Luther SA. CCL21 is sufficient to mediate DC migration, maturation and function in the absence of CCL19. European Journal of Immunology (2010) 40:1266–1271. doi: 10.1002/eji.200939921

40. Scandella E, Men Y, Legler DF, Gillessen S, Prikler L, Ludewig B, Groettrup M. CCL19/CCL21-triggered signal transduction and migration of dendritic cells requires prostaglandin E2. Blood (2004) 103:1595–1601. doi: 10.1182/blood-2003-05-1643

41. Joutoku Z, Onodera T, Matsuoka M, Homan K, Momma D, Baba R, Hontani K, Hamasaki M, Matsubara S, Hishimura R, et al. CCL21/CCR7 axis regulating juvenile cartilage repair can enhance cartilage healing in adults. Scientific Reports (2019) 9:1–12. doi: 10.1038/s41598-019-41621-3

42. Li F, Zou Z, Suo N, Zhang Z, Wan F, Zhong G, Qu Y, Ntaka KS, Tian H. CCL21/CCR7 axis activating chemotaxis accompanied with epithelial–mesenchymal transition in human breast carcinoma. Medical Oncology (2014) 31:1–10. doi: 10.1007/s12032-014-0180-8

43. Sasaki M, Abe R, Fujita Y, Ando S, Inokuma D, Shimizu H. Mesenchymal Stem Cells Are Recruited into Wounded Skin and Contribute to Wound Repair by Transdifferentiation into Multiple Skin Cell Type. The Journal of Immunology (2008) 180:2581–2587. doi: 10.4049/jimmunol.180.4.2581

44. Mo M, Zhou M, Wang L, Qi L, Zhou K, Liu LF, Chen Z, Zu XB. CCL21/CCR7 enhances the proliferation, migration, and invasion of human bladder cancer T24 cells. PLoS ONE (2015) 10:1–12. doi: 10.1371/journal.pone.0119506

45. Xiong Y, Huang F, Li X, Chen Z, Feng D, Jiang H, Chen W, Zhang X. CCL21/CCR7 interaction promotes cellular migration and invasion via modulation of the MEK/ERK1/2 signaling pathway and correlates with lymphatic metastatic spread and poor prognosis in urinary bladder cancer. International Journal of Oncology (2017) 51:75–90. doi: 10.3892/ijo.2017.4003

46. Aberle H. Axon guidance and collective cell migration by substrate-derived attractants. Frontiers in Molecular Neuroscience (2019) 12:1–7. doi: 10.3389/fnmol.2019.00148

47. Cooper JA. Mechanisms of cell migration in the nervous system. Journal of Cell Biology (2013) 202:725–734. doi: 10.1083/jcb.201305021

48. Borrell V, Marín O. Meninges control tangential migration of hem-derived Cajal-Retzius cells via CXCL12/CXCR4 signaling. Nature Neuroscience (2006) 9:1284–1293. doi: 10.1038/nn1764

49. Doitsidou M, Reichman-Fried M, Stebler J, Köprunner M, Dörries J, Meyer D, Esguerra C v, Leung TC, Raz E. Guidance of primordial germ cell migration by the chemokine SDF-1. Cell (2002) 111:647–659. doi: 10.1016/S0092-8674(02)01135-2

50. Opatz J, Küry P, Schiwy N, Järve A, Estrada V, Brazda N, Bosse F, Müller HW. SDF-1 stimulates neurite growth on inhibitory CNS myelin. Molecular and Cellular Neuroscience (2009) doi: 10.1016/j.mcn.2008.11.002

51. Pujol F, Kitabgi P, Boudin H. The chemokine SDF-1 differentially regulates axonal elongation and branching in hippocampal neurons. Journal of Cell Science (2005) 118:1071–1080. doi: 10.1242/jcs.01694

52. Hirth M, Gandla J, Höper C, Gaida MM, Agarwal N, Simonetti M, Demir A, Xie Y, Weiss C, Michalski CW, et al. CXCL10 and CCL21 Promote Migration of Pancreatic Cancer Cells Toward Sensory Neurons and Neural Remodeling in Tumors in Mice, Associated With Pain in Patients. Gastroenterology (2020) 159:665–681.e13. doi: 10.1053/j.gastro.2020.04.037

53. Liu JX, Cao X, Tang YC, Liu Y, Tang FR. CCR7, CCR8, CCR9 and CCR10 in the mouse hippocampal CA1 area and the dentate gyrus during and after pilocarpine-induced status epilepticus. Journal of Neurochemistry (2007) 100:1072–1088. doi: 10.1111/j.1471-4159.2006.04272.x

54. Comerford I, Harata-Lee Y, Bunting MD, Gregor C, Kara EE, McColl SR. A myriad of functions and complex regulation of the CCR7/CCL19/CCL21 chemokine axis in the adaptive immune system. Cytokine and Growth Factor Reviews (2013) 24:269–283. doi: 10.1016/j.cytogfr.2013.03.001

55. Shannon LA, Calloway PA, Welch TP, Vines CM. CCR7/CCL21 migration on fibronectin is mediated by phospholipase Cγ1 and ERK1/2 in primary T lymphocytes. Journal of Biological Chemistry (2010) 285:38781–38787. doi: 10.1074/jbc.M110.152173

56. Hauser MA, Legler DF. Common and biased signaling pathways of the chemokine receptor CCR7 elicited by its ligands CCL19 and CCL21 in leukocytes. Journal of Leukocyte Biology (2016) 99:869–882. doi: 10.1189/jlb.2mr0815-380r

57. Jørgensen AS, Rosenkilde MM, Hjortø GM. Biased signaling of G protein-coupled receptors – From a chemokine receptor CCR7 perspective. General and Comparative Endocrinology (2018) 258:4–14. doi: 10.1016/j.ygcen.2017.07.004

58. Hjortø GM, Larsen O, Steen A, Daugvilaite V, Berg C, Fares S, Hansen M, Ali S, Rosenkilde MM. Differential CCR7 targeting in dendritic cells by three naturally occurring CC-chemokines. Frontiers in Immunology (2016) 7:1–15. doi: 10.3389/fimmu.2016.00568

59. Corbisier J, Galès CL, Huszagh A, Parmentier M, Springael JY. Biased signaling at chemokine receptors. Journal of Biological Chemistry (2015) 290:9542–9554. doi: 10.1074/jbc.M114.596098

60. Kohout TA, Nicholas SL, Perry SJ, Reinhart G, Junger S, Struthers RS. Differential desensitization, receptor phosphorylation, β-arrestin recruitment, and ERK1/2 activation by the two endogenous ligands for the CC chemokine receptor 7. Journal of Biological Chemistry (2004) 279:23214–23222. doi: 10.1074/jbc.M402125200

61. Shannon LA, McBurney TM, Wells MA, Roth ME, Calloway PA, Bill CA, Islam S, Vines CM. CCR7/CCL19 controls expression of EDG-1 in T cells. Journal of Biological Chemistry (2012) 287:11656–11664. doi: 10.1074/jbc.M111.310045

62. Riol-Blanco L, Sánchez-Sánchez N, Torres A, Tejedor A, Narumiya S, Corbí AL, Sánchez-Mateos P, Rodríguez-Fernández JL. The Chemokine Receptor CCR7 Activates in Dendritic Cells Two Signaling Modules That Independently Regulate Chemotaxis and Migratory Speed. The Journal of Immunology (2005) 174:4070–4080. doi: 10.4049/jimmunol.174.7.4070

63. Cai D, Yingjing S, de Bellard ME, Tang S, Filbin MT. Prior exposure to neurotrophins blocks inhibition of axonal regeneration by MAG and myelin via a cAMP-dependent mechanism. Neuron (1999) 22: doi: 10.1016/S0896-6273(00)80681-9

64. Perlson E, Hanz S, Ben-Yaakov K, Segal-Ruder Y, Seger R, Fainzilber M. Vimentin-dependent spatial translocation of an activated MAP kinase in injured nerve. Neuron (2005) 45: doi: 10.1016/j.neuron.2005.01.023

65. Chierzi S, Ratto GM, Verma P, Fawcett JW. The ability of axons to regenerate their growth cones depends on axonal type and age, and is regulated by calcium, cAMP and ERK. European Journal of Neuroscience (2005) 21: doi: 10.1111/j.1460-9568.2005.04066.x

66. Puttagunta R, Tedeschi A, Sória MG, Hervera A, Lindner R, Rathore KI, Gaub P, Joshi Y, Nguyen T, Schmandke A, et al. PCAF-dependent epigenetic changes promote axonal regeneration in the central nervous system. Nature Communications (2014) 5: doi: 10.1038/ncomms4527

67. Hanz S, Fainzilber M. Retrograde signaling in injured nerve - The axon reaction revisited. Journal of Neurochemistry (2006) 99: doi: 10.1111/j.1471-4159.2006.04089.x

68. Leite SC, Pinto-Costa R, Sousa MM. Actin dynamics in the growth cone: a key player in axon regeneration. Current Opinion in Neurobiology (2021) 69:11–18. doi: 10.1016/j.conb.2020.11.015

69. Tanimura S, Takeda K. ERK signalling as a regulator of cell motility. Journal of Biochemistry (2017) 162:145–154. doi: 10.1093/jb/mvx048

70. Danson CM, Pocha SM, Bloomberg GB, Cory GO. Phosphorylation of WAVE2 by MAP kinases regulates persistent cell migration and polarity. Journal of Cell Science (2007) 120:4144–4154. doi: 10.1242/jcs.013714

71. Martinez-Quiles N, Ho H-YH, Kirschner MW, Ramesh N, Geha RS. Erk/Src Phosphorylation of Cortactin Acts as a Switch On-Switch Off Mechanism That Controls Its Ability To Activate N-WASP. Molecular and Cellular Biology (2004) 24:5269–5280. doi: 10.1128/mcb.24.12.5269-5280.2004

72. Mendoza MC, Er EE, Zhang W, Ballif BA, Elliott HL, Danuser G, Blenis J. ERK-MAPK Drives Lamellipodia Protrusion by Activating the WAVE2 Regulatory Complex. Molecular Cell (2011) 41:661–671. doi: 10.1016/j.molcel.2011.02.031

73. Nakanishi O, Suetsugu S, Yamazaki D, Takenawa T. Effect of WAVE2 phosphorylation on activation of the Arp2/3 complex. Journal of Biochemistry (2007) 141:319–325. doi: 10.1093/jb/mvm034

74. Tong J, Li L, Ballermann B, Wang Z. Phosphorylation of Rac1 T108 by Extracellular Signal-Regulated Kinase in Response to Epidermal Growth Factor: a Novel Mechanism To Regulate Rac1 Function. Molecular and Cellular Biology (2013) 33:4538–4551. doi: 10.1128/mcb.00822-13

75. Pollard TD. Actin and actin-binding proteins. Cold Spring Harbor Perspectives in Biology (2016) 8:1–17. doi: 10.1101/cshperspect.a018226

76. Ruiz-Miguel JES, Letourneau PC. The role of Arp2/3 in growth cone actin dynamics and guidance is substrate dependent. Journal of Neuroscience (2014) 34:5895–5908. doi: 10.1523/JNEUROSCI.0672-14.2014

77. Schlau M, Terheyden-Keighley D, Theis V, Mannherz HG, Theiss C. VEGF triggers the activation of cofilin and the Arp2/3 complex within the growth cone. International Journal of Molecular Sciences (2018) 19: doi: 10.3390/ijms19020384

78. Nossent AY, Bastiaansen AJNM, Peters EAB, de Vries MR, Aref Z, Welten SMJ, de Jager SCA, van der Pouw Kraan TCTM, Quax PHA. CCR7-CCL19/CCL21 axis is essential for effective arteriogenesis in a murine model of hindlimb ischemia. Journal of the American Heart Association (2017) 6:1–17. doi: 10.1161/JAHA.116.005281

79. Herrmann KA, Broihier HT. What neurons tell themselves: autocrine signals play essential roles in neuronal development and function. Current Opinion in Neurobiology (2018) 51:70–79. doi: 10.1016/j.conb.2018.03.002

80. Acheson A, Conover JC, Fandl JP, DeChiara TM, Russell M, Thadani A, Squinto SP, Yancopoulos GD, Lindsay RM. A BDNF autocrine loop in adult sensory neurons prevents cell death. Nature (London) (1995) 374:450–453. doi: 10.1038/374450a0

81. Cheng PL, Song AH, Wong YH, Wang S, Zhang X, Poo MM. Self-amplifying autocrine actions of BDNF in axon development. Proceedings of the National Academy of Sciences of the United States of America (2011) 108:18430–18435. doi: 10.1073/pnas.1115907108

82. Froger N, Matonti F, Roubeix C, Forster V, Ivkovic I, Brunel N, Baudouin C, Sahel JA, Picaud S. VEGF is an autocrine/paracrine neuroprotective factor for injured retinal ganglion neurons. Scientific Reports (2020) 10:1–11. doi: 10.1038/s41598-020-68488-z

83. Ogunshola OO, Antic A, Donoghue MJ, Fan SY, Kim H, Stewart WB, Madri JA, Ment LR. Paracrine and autocrine functions of neuronal vascular endothelial growth factor (VEGF) in the central nervous system. Journal of Biological Chemistry (2002) 277:11410–11415. doi: 10.1074/jbc.M111085200

84. Baliki MN, Apkarian AV. Nociception, Pain, Negative Moods, and Behavior Selection. Neuron (2015) 87:474–491. doi: 10.1016/j.neuron.2015.06.005

85. Woller SA, Eddinger KA, Corr M, Yaksh TL, Corr M, Yaksh TL. Woller S.A, Eddinger K.A, Corr M, Yaksh T.L, An overview of pathways encoding nociception. Clin Exp Rheumatol 2017; 35 (suppl. 107) S40–S46. (2017)

86. Rabiller L, Labit E, Guissard C, Gilardi S, Guiard BP, Moulédous L, Silva M, Mithieux G, Pénicaud L, Lorsignol A, et al. Pain sensing neurons promote tissue regeneration in adult mice. npj Regenerative Medicine (2021) 6:1–9. doi: 10.1038/s41536-021-00175-7

87. Amadesi S, Reni C, Katare R, Meloni M, Oikawa A, Beltrami AP, Avolio E, Cesselli D, Fortunato O, Spinetti G, et al. Role for substance P-based nociceptive signaling in progenitor cell activation and angiogenesis during ischemia in mice and in human subjects. Circulation (2012) 125:1774–1786. doi: 10.1161/CIRCULATIONAHA.111.089763

88. Wei JJ, Kim HS, Spencer CA, Brennan-Crispi D, Zheng Y, Johnson NM, Rosenbach M, Miller C, Leung DH, Cotsarelis G, et al. Activation of TRPA1 nociceptor promotes systemic adult mammalian skin regeneration. Science Immunology (2020) 5:1–9. doi: 10.1126/SCIIMMUNOL.ABA5683

89. Donnelly DJ, Popovich PG. Inflammation and its role in neuroprotection, axonal regeneration and functional recovery after spinal cord injury. Experimental Neurology (2008) doi: 10.1016/j.expneurol.2007.06.009

90. Hervera A, de Virgiliis F, Palmisano I, Zhou L, Tantardini E, Kong G, Hutson T, Danzi MC, Perry RBT, Santos CXC, et al. Reactive oxygen species regulate axonal regeneration through the release of exosomal NADPH oxidase 2 complexes into injured axons. Nature Cell Biology (2018) 20:307–319. doi: 10.1038/s41556-018-0039-x

