## Supplementary figures and images for "Nociception-dependent CCL21 induce dorsal root ganglia axonal growth via CCR7-ERK activation"

### Supplementary Figure 1

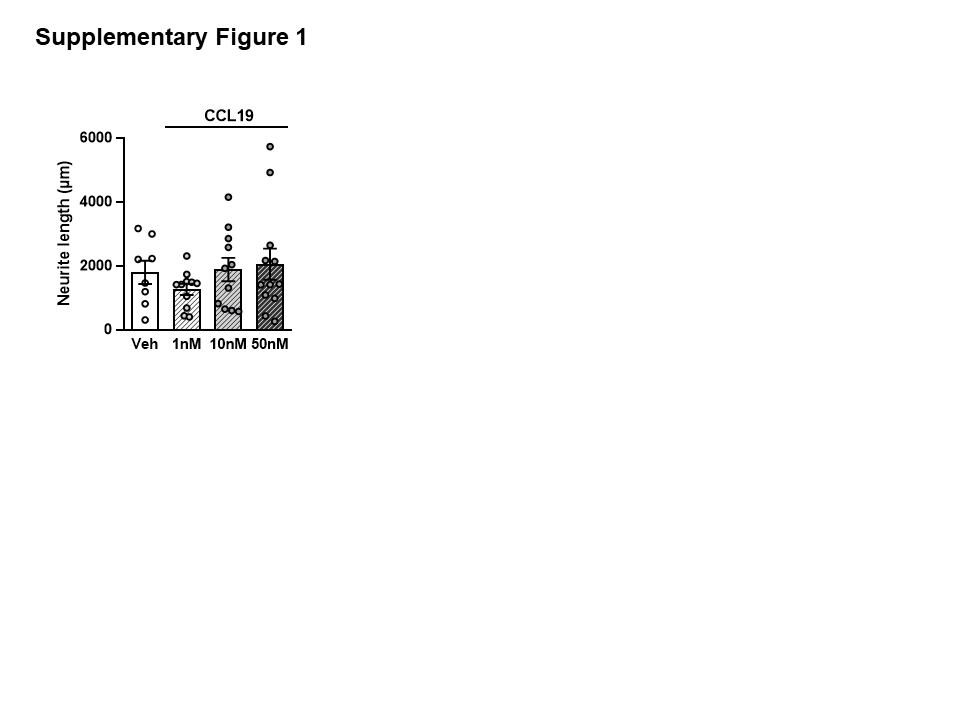

### Supplementary Figure 2

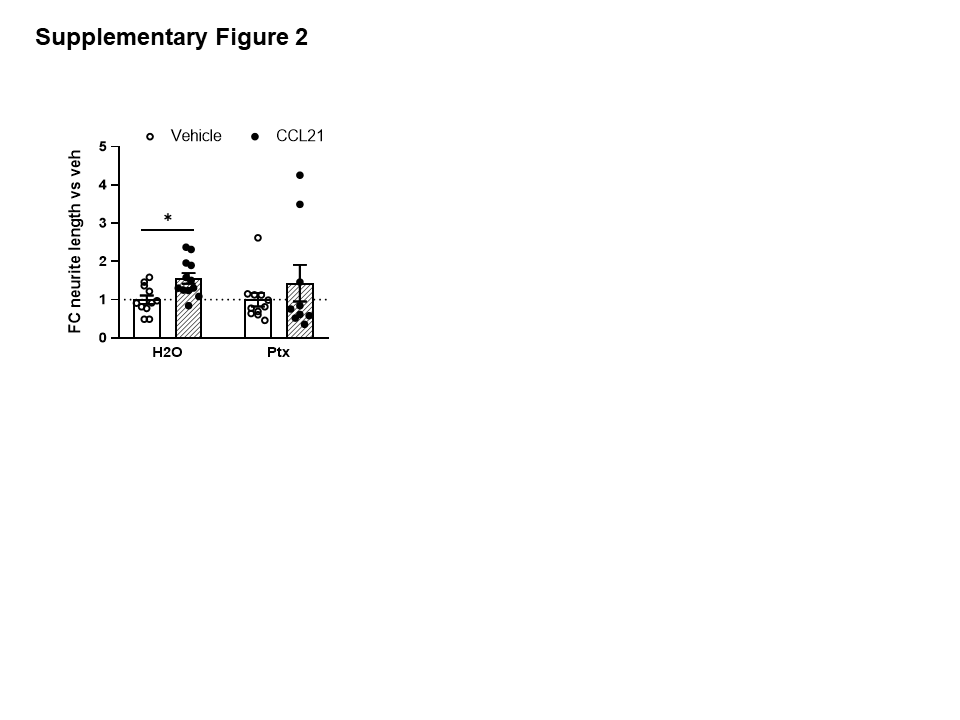
